# Anti-CoVid19 plasmid DNA vaccine induces a potent immune response in rodents by Pyro-drive Jet Injector intradermal inoculation

**DOI:** 10.1101/2021.01.13.426436

**Authors:** Tomoyuki Nishikawa, Chin Yang Chang, Jiayu A Tai, Hiroki Hayashi, Jiao Sun, Shiho Torii, Chikako Ono, Yoshiharu Matsuura, Ryoko Ide, Junichi Mineno, Miwa Sasai, Masahiro Yamamoto, Hironori Nakagami, Kunihiko Yamashita

## Abstract

There is an urgent need to limit and stop the worldwide coronavirus disease 2019 (COVID-19) pandemic via quick development of efficient and safe vaccination methods. Plasmid DNA vaccines are one of the most remarkable vaccines that can be developed in a short term. pVAX1-SARS-CoV2-co, which is a plasmid DNA vaccine, was designed to express severe acute respiratory syndrome coronavirus-2 (SARS-CoV-2) spike protein. The produced antibodies lead to Immunoreactions against S protein, anti-receptor-binding-domain, and neutralizing action of pVAX1-SARS-CoV2-co, as confirmed in a previous study. To promote the efficacy of the pVAX1-SARS-CoV2-co vaccine, a pyro-drive jet injector (PJI) was employed. PJI is an injection device that can adjust the injection pressure depending on various target tissues. Intradermally-adjusted PJI demonstrated that pVAX1-SARS-CoV2-co vaccine injection caused a strong production of anti-S protein antibodies, triggered immunoreactions and neutralizing actions against SARS-CoV-2. Moreover, a high dose of pVAX1-SARS-CoV2-co intradermal injection via PJI did not cause any serious disorders in the rat model. Finally, virus infection challenge in mice, confirmed that intradermally immunized (via PJI) mice were potently protected from COVID-19 infection. Thus, pVAX1-SARS-CoV2-co intradermal injection via PJI is a safe and promising vaccination method to overcome the COVID-19 pandemic.

## Introduction

The coronavirus disease 2019 (COVID-19) pandemic accelerated the development of a safe and effective vaccine candidate against the severe acute respiratory syndrome coronavirus-2 (SARS-CoV-2). At least 52 vaccine candidates have been tested in clinical studies, and several types of vaccine platforms were used, such as protein subunit, replicating viral vector, non-replicating viral vector, inactivated virus, RNA, DNA, live attenuated virus, and virus-like particles (VLP); the number of each vaccine candidate was 16, 5, 9, 7, 6, 6,1, and 2, respectively. Furthermore, an intramuscular injection was employed for 44 vaccine candidates and an intradermal injection method was employed for three DNA vaccine candidates and other five vaccine candidates were administrated subcutaneously, intranasally, or via oral route (Table 1)^1^. Plasmid DNA^2^ and mRNA^3^ vaccines have been developed in addition to traditional vaccines because both can be quickly produced via generic manufacturing processes and can be constructed directly from the genetic sequence information. This means that the nucleotide-based vaccines are easy to adapt to viral mutations^2^. Furthermore, mass production technology has been established for DNA plasmids and new technologies are being developed for mRNA vaccines as well, such as mRNA stabilizing technology^3^. Five mRNA vaccine candidates and four DNA vaccine candidates are involved in developing a vaccine against SARS-CoV-2 virus. In contrast, an effective method to introduce these molecules, especially DNA plasmids, into cells should be established for effective protein expression and antibody induction. To resolve this issue, we previously reported the potential of the intradermal jet injection method for DNA vaccination via model DNA vaccination using ovalbumin expression plasmid^4^. In addition, when developing a new DNA vaccine, it is important to assess the health effect of new vaccine candidates in addition to efficacy evaluation.

**Table 1.**
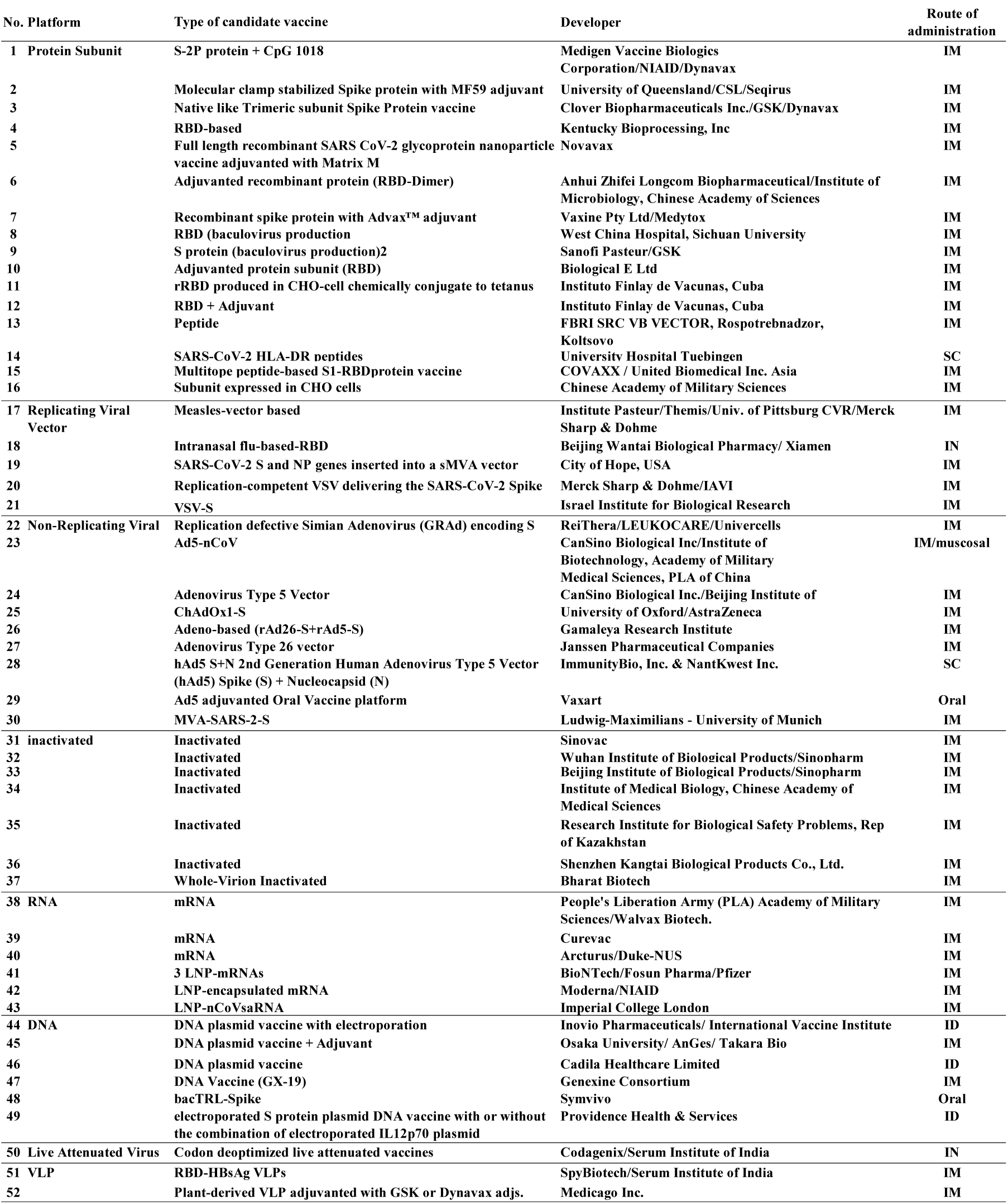
List of COVID-19 vaccine candidates

Nucleic acid-based vaccines have a great advantage of being developed at an accelerated pace^5^. In addition, DNA vaccines are thermo-stable, whereas RNA vaccines require a cold-chain system, which can be an obstacle for the worldwide supply of anti-COVID-19 vaccines. Recently, novel plasmid DNA vaccines (AG0302-COVID19) against SARS-CoV-2 virus have been developed and are currently in phase I/II trials in Japan^6^. AG0302-COVID19 has an optimized S protein sequence that accelerates S protein expression in human cells. The S protein expression in AG0302-COVID-transfected cells was successfully confirmed. AG0302-COVID19 with alum adjuvant was intramuscularly injected, which induced both humoral and cellular immune responses in a preclinical study. Moreover, AG0302-COVID19-induced antibodies showed neutralizing activity for SARSCoV-2 S protein binding. Results of the peptide array used to analyze the antibody epitopes revealed that most AG0302-COVID19-induced antibodies recognized S2 and receptor binding domain (RBD). The anti-COVID-19 DNA vaccine candidate pVAX1-SARS-CoV2-co has the same nucleotide sequence as AG0302-COVID19. The pVAX1-SARS-CoV2-co-triggered antibodies have also been confirmed to neutralize RBD recombinant protein and angiotensin-converting enzyme 2 (ACE2), which is a SARS-CoV-2 receptor.

We employed a novel pyro-drive jet injector (PJI) was to enhance the pVAX1-SARS-CoV2-co efficacy via dermal injection while previously, an intramuscular injection was used. One of the limitations of DNA vaccines is the low antibody titer against COVID-19 infections^5^. PJI can be applied for injecting pVAX1-SARS-CoV2-co into dermal tissues, where various immune cells are stimulated upon S protein expression. In PJI, combustion of two types of explosives leads to the plunger discharging liquid samples in the container toward the target tissue at very high pressure^7^. The great advantage of PJI is the adjustment of sample dose and depth of delivery, which can be targeted by changing the two types of explosives. Different plasmid DNA, luciferase and ovalbumin (OVA) were successfully delivered to dermal tissue and much higher expression of these proteins was detected in the target tissue than the classical needle syringe injection.

Moreover, the antibody titer against OVA injection by PJI is more than a hundred times higher than needle syringe injection at 8 weeks after OVA plasmid DNA injection^4^. This study aimed to determine the safety of a new anti-SARS-CoV-2 DNA vaccine candidate via intradermal jet injection. Therefore, the adjusted PJI to deliver pVAX1-SARS-CoV2-co to dermal tissue was employed, and potent anti-COVID-19 immune reactions were confirmed in this study.

## Results

### pVAX-SARS-CoV2-co vaccine injection to mouse and rat dermal tissue via PJI

A novel injection device, PJI, can introduce pVAX1-SARS-CoV2-co to mouse dermal tissue. Exactly 20 or 50 μl of the pVAX1-SARS-CoV2-co solution was injected into the mouse or rat back using PJI. An approximately 5-mm wheal in diameter was formed on the back of the mouse/rat and no bleeding or inflammation observed (Fig1, A, B). Therefore, pVAX1-SARS-CoV2-co could be efficiently introduced to rat dermal tissue via PJI.

**Figure 1.**
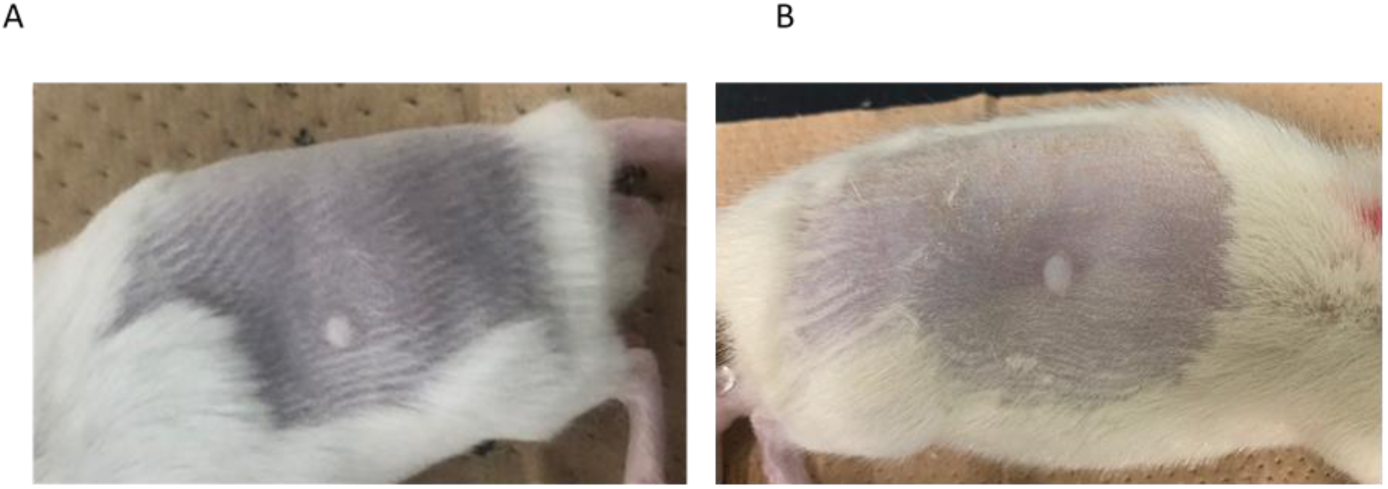
pVAX1-SARS-CoV2-co injection to dermal tissue by Pyro-drive jet injector. A) A wheal on the back of the rat right after 20 μl injection using PJI. B) A wheal on rat back right after 50 μl injection by PJI.

### The increase and retention of anti-COVID19 spike protein antibody in the rat serum

pVAX1-SARS-CoV2-co was injected into the back of the rats three times every other week using PJI (Fig 2A). Blood samples were collected every other week until 12 weeks after the first injection, and sera were purified from the blood samples. Anti-COVID19 spike protein (S1+S2) antibody titer at 2, 4, and 6 weeks after the first injection was measured using enzyme-linked immunosorbent assay (ELISA). After 2 weeks, the antibody titer started to rise and reached the highest value after 6 weeks (Fig 2B). The antibody titer was measured until after 12 weeks to confirm antibody retention upon pVAX1-SARS-CoV2-co injection. The antibody titer was the highest after 6 weeks and gradually decreased toward week 12, but all titers were still very high until 12 weeks (Fig 2C). To determine the polarization of immunoglobulin subclass profile in rat sera following vaccine treatment, we analyzed the differences in SARS-CoV2-specific IgG1, IgG2a, IgG2b, and IgG2c in rat sera harvested 8 weeks after vaccine treatment. Our results showed that overall SARS-CoV2-specific IgG1, IgG2a, and IgG2b levels were high, whereas IgG2c levels were low (Fig 2D). Importantly, the SARS-CoV2-specific IgG2b subclass was more elevated than IgG1, indicating that Th1 mediated subclass polarization. Additionally, antibody titers for RBD of the spike glycoprotein in rat serum were measured. The RBD titers were elevated weeks 4 and 12 after the first vaccination (Fig 2E). Production of both anti-S1+S2 glycoprotein and anti-RBD antibodies indicated that the intradermal pVAX1-SARS-CoV2-co injection successfully induced an immune response.

**Figure 2.**
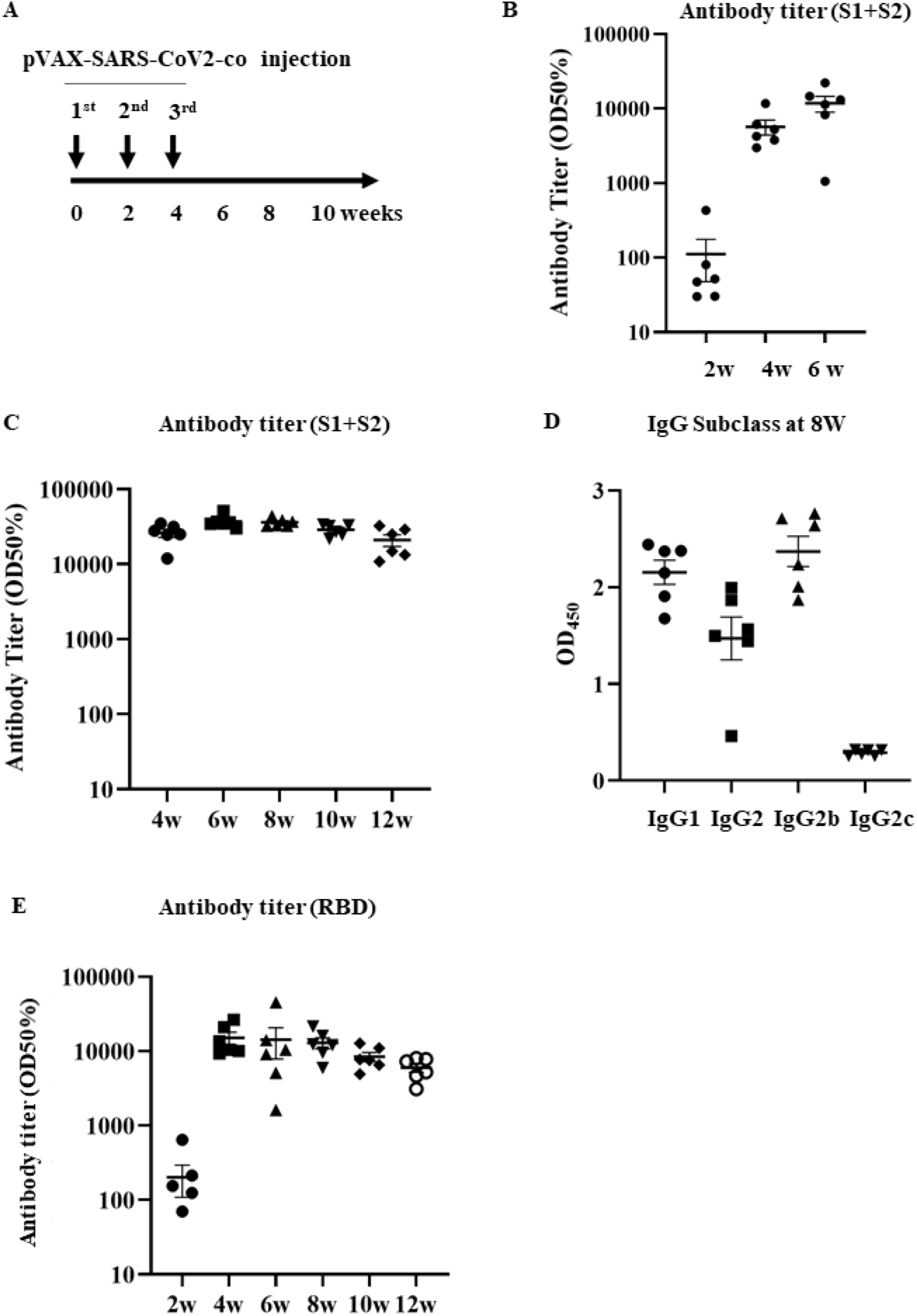
pVAX1-SARS-CoV2-co vaccination animal protocol and humoral immune responses. A) Vaccination protocol. pVAX1-SARS-CoV2-co was injected intradermally using PJI three times at 2-week intervals. Blood samples were collected every two weeks for antibody titer and subclass analysis. B) Early (2 to 6 weeks after the first injection) antibody titer (half-maximum) for recombinant glycoprotein S1+S2 assessed using ELISA. C) Antibody titer (half-maximum) for recombinant glycoprotein S1+S2 (4 to 12 weeks) D) Immunoglobulin subclasses of rat serum 8 weeks after the vaccine injection. IgG subclasses in rat sera (IgG1, IgG2a, IgG2b and IgG2c) were analyzed using ELISA. The results were assessed at 450 nm. E) Antibody titer (half-maximum) for recombinant spike glycoprotein RBD assessed using ELISA (2 to 12 weeks).

### The cellular immune response of rats upon vaccination

To examine Th1/Th2 cellular immune responses to the vaccine, we isolated solenocytes from vaccinated rats 2 weeks after the final vaccination and analyzed for S1+S2-specific IFN-γ and IL-4-secreting T cells using ELISpot assay. Splenocytes harvested from vaccinated rats showed a significant increase in IFN-γ-secreting S1+S2-specific T cells (Fig. 3A), whereas IL-4-secreting S1+S2-specific T cells were detected at much lower numbers (data not shown). This indicated that intradermal injection of pVAX1-SARS-CoV2-co induced strong Th1-type cellular immune response.

**Figure 3.**
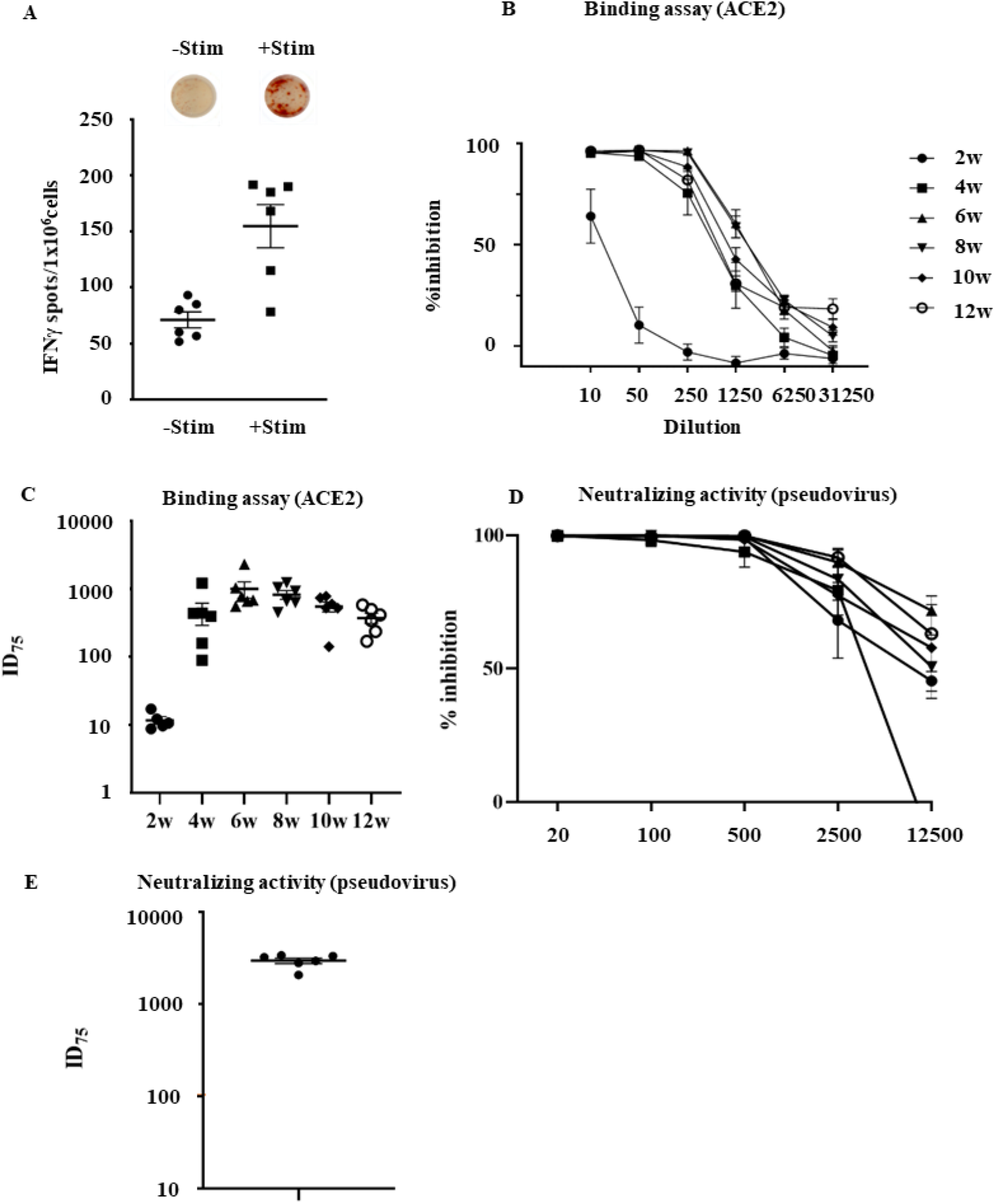
A cellular immune response and neutralizing antibodies. A) IFN-γ and IL-4 ELISpot responses in splenocytes from rats vaccinated with 240 μg pVAX1-SARS-CoV2-co, with or without re-stimulation with recombinant S1+S2. The p-value calculated using Welch’s t-test between IFN-γ secreting cell number in the antigen-stimulated group and the un-stimulated group was 0.0059 (n=6). B) ACE2 (and S1+S2) binding inhibition assay using 2-to 12-week immunized rat serum normalized to pre-serum, at reciprocal 10-to 31250-fold dilutions. C) Neutralization titers (ID75) of 2-to 12-week immunized rat serum indicated as the dilution of serum required for 75% inhibition of ACE2-S1+S2 binding as shown in (B). D) Inhibition rate against the pseudovirus. The immunized rat serum at 8 weeks was used at reciprocal 20-to 12500-fold dilutions. E) Neutralization activity (ID75; 75% inhibition). The values indicate the inhibitory dose of the serum which shows 75% inhibition rate against pseudovirus binding as shown in (D).

### ACE2 binding inhibition and pseudovirus neutralization assays

To further confirm the neutralizing activity of vaccine-induced antibodies against SARS-CoV-2, we analyzed the binding inhibition of human ACE2, a receptor of SARS-Cov-2 spike glycoprotein, with S1+S2 recombinant protein with immunized rat serum using ELISA^8^. Rat serum collected 2 weeks after the first vaccination showed some inhibition ability (62%) against hACE2 binding of to the S1+S2 recombinant protein at a 10-fold dilution (Fig 3D). In comparison, rat serum collected between weeks 4 and 12 showed high neutralizing ability (>90%) against the binding of hACE2 to S1+S2 recombinant protein at up to 50 folds dilution and maintained high neutralization activity for 6-8 weeks at up to 250 folds dilution. Correspondingly, neutralization titers against hACE2-S1+S2 binding (at 75% inhibitory/neutralization dose (ID75)) showed a significant jump between weeks 2 and 4 after the first vaccination, reaching a peak between 6 and 8 weeks, then decreasing gradually after 8 weeks (Fig 3E). Moreover, pseudovirus neutralization assay, using pseudotyped vesicular stomatitis virus (VSV) with the luciferase gene and Vero E6 cells stably expressing TMPRSS2^9-10^, was performed to examine the neutralizing activity. A series of rat serum dilutions 8 weeks after the first vaccination showed strong neutralizing activity against pseudovirus infection (Fig 3D). Neutralizing titers (ID75) exhibited the highest titer after 6 weeks, which was about thirty times higher than those after intramuscular injection(Fig 3E).

### Increasing injection doses

To assess whether vaccine dosage would affect antibody production and cellular immune responses, we assessed the pVAX1-SARS-CoV2-co dosage in the range of 100 (low dose) to 400 µg (high dose). When 100 or 400 µg of DNA vaccine was injected into the rat skin, the antibody production was increased from week 2 to 6 after every injection in both doses. The maximum antibody titers of the 100 and 400µg DNA vaccination groups were 17056±6331 or 59068±33578 after 6 weeks, respectively. In addition, there was a statistically significant difference between the antibody titers of 100 and 400 µg injected group after both 4 and 6 weeks (Fig 4A). To assess whether vaccine dosage would affect Th1/Th2 cellular immune responses, we compared S1+S2-specific IFN-γ and IL-4 ELISpot assays between rats vaccinated with 100 μg or 400 μg of pVAX1-SARS-CoV2-co. Splenocytes harvested from 400 μg/high-dose vaccinated rats showed a significant increase in S1+S2-specific IFN-γ secretion when re-stimulated with the antigen, but those from low-dose vaccinated rats exhibited only a modest increase when re-stimulated (Fig 4B). IL-4-secreting T cells were barely detected in both vaccinated groups. These results suggested that a high-dose vaccine could induce a stronger IFN-γ response compared to the lower dose vaccine, which could lead to a more predominant Th1-driven cellular immune response.

**Figure 4.**
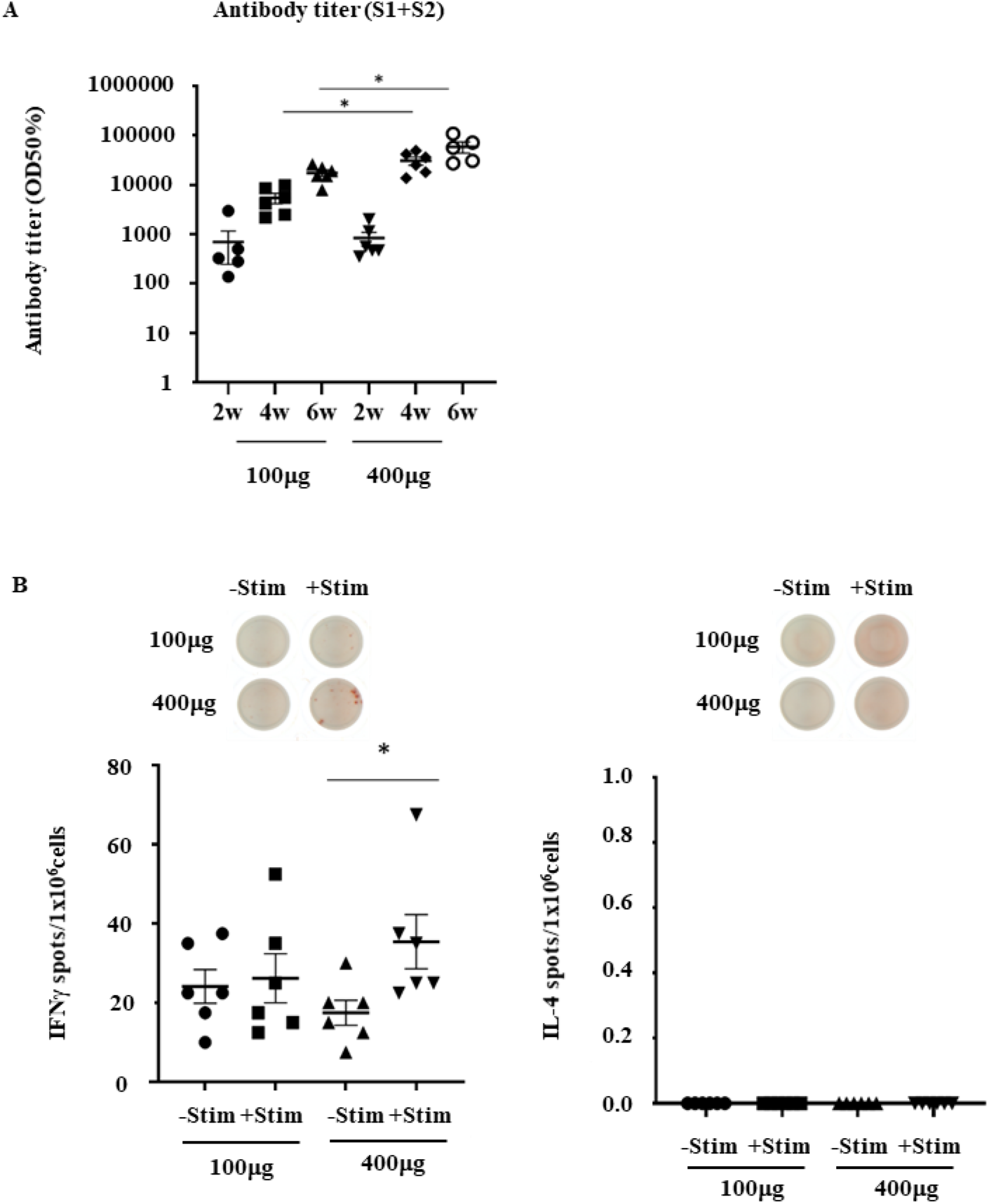
Increasing injection doses; 100 μg (low dose) and 400 μg (high dose). A) Antibody titer (half-maximum) for recombinant glycoprotein S1+S2 (2 to 6 weeks) of 100 and 400 μg of pVAX1-SARS-CoV2-co-injected group. The exact p-values calculated using Welch’s t-test between 100 and 400 µg doses were 0.0060 or 0.0496 after 4 and 6 weeks, respectively. Both p-values were less than 0.05. These results indicated that a statistically significant difference was observed between 100- and 400-µg doses. B) IFN-γ and IL-4 ELISpot responses in splenocytes from rats vaccinated with 100 or 400 μg of pVAX1-SARS-CoV2-co-injected group, with or without re-stimulation with recombinant S1+S2. The exact p-values calculated using Student’s *t*-test between -Stim and +Stim was *P*=0.0394.

### DNA vaccine prevents viral infection challenge in mice

To assess whether DNA vaccination could prevent viral infection, we injected mice with the mouse-adapted SARS-CoV-2 virus (HuDPKng19-020 strain - Q498Y/P499T)^11^ intradermally (ID; PJI). Prior to infection challenge 16 weeks post immunization, the antibody titers against S1+S2 in the ID injection group were significantly higher than those in the non-vaccinated (NV) group (Fig. 5A, Supplemental Fig. 1A). Similarly, the antibody titers for RBD in the ID injection group were higher than those in the NV group (Fig. 5B, Supplemental Fig. 1B). ID injection group also displayed stronger inhibitory ability and higher neutralization titers against hACE2-S1+S2 binding than the NV group (Fig. 5C). After virus infection challenge, no detectable viral load was found in the lungs of all eight mice in the ID injection group (Fig. 5D, Supplemental Fig. 1D). Taken together, these results suggested that pVAX1-SARS-CoV2-co immunized mice, especially mice vaccinated via PJI, could effectively achieve upregulated antibody responses against the spike protein of SARS-CoV-2.

**Figure 5.**
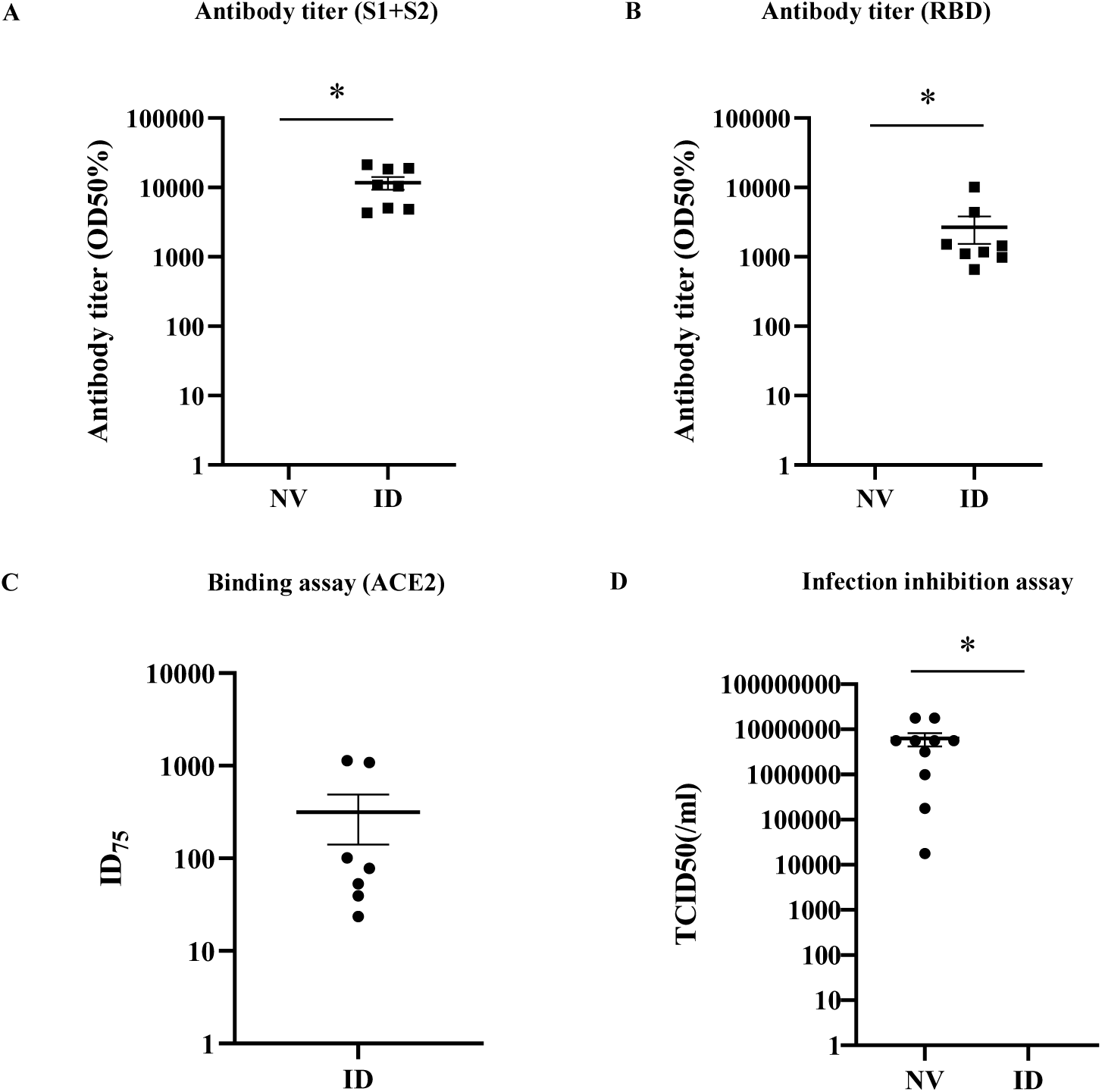
Virus challenge on pVAX1-SARS-CoV2-co immunized mice A) Pre-viral challenge of mice immunized with 160 μg of pVAX1-SARS-CoV2-co plasmid intradermally (ID; n = 8) or no-vaccinated (NV: n=10) two times at 2-week intervals. Antibody titer (half maximum) for recombinant S1+S2 in the blood serum 16 weeks post initial vaccination, assessed using ELISA. *P*<0.001 Student’s *t*-test. B) Antibody titer (half maximum) for recombinant RBD in the blood serum in ID and NV mice. *P*=0.018 Student’s *t*-test. C) Neutralization titer (ID75) for recombinant ACE2-S1+S2 binding inhibition in the blood serum of ID mice. D) ID and NV (n=10) mice were intranasally infected with mouse-adapted SARS-CoV-2 virus. 50% tissue culture infective doses (TCID50) values in the lung tissues of each animal are shown. *P*=0.016 Student’s *t*-test. All individual values are shown in Supplemental Fig. 1.

### Safety assessment for pVAX1-SARS-CoV2-co intradermal injection and the immune response

No animals had died at the time of necropsy or showed abnormalities except for skin reaction. There was no significant difference in body weight, food consumption, and hematology between the PBS-injected groups and DNA vaccination groups (Tables 2 to 4). In addition, the urinalysis data did not show any toxic signs (data not shown). As shown in Table 5, blood biochemistry revealed significantly lower levels of triglyceride and β-globulin in the 400 µg group compared to the PBS group. After DNA vaccination, erythema and scab formations were observed at the injection site for the rats injected with the vaccine, whereas no erythema or scab formation was observed in the PBS-injected group (Table 6). In the 100-µg DNA vaccine injected group, scab formation occurred 5, 4, or 3 days after the first, second, or third vaccination, respectively. In contrast, in the 400-μg DNA vaccine injected group, scab formation occurred 5, 3, or 2 days after the first, second, or third vaccination, respectively. The scab formation lasted shorter in the high dose than that in the low-dose group. In addition, recovery from the scab formation was prolonged depending on the vaccination time. After the first, second, and third injections, the scab disappeared after 3, 5–7, and 11 days, respectively. However, all scabs disappeared and the skin recovered 14 days after every injection (Table 6).

**Table 2.**
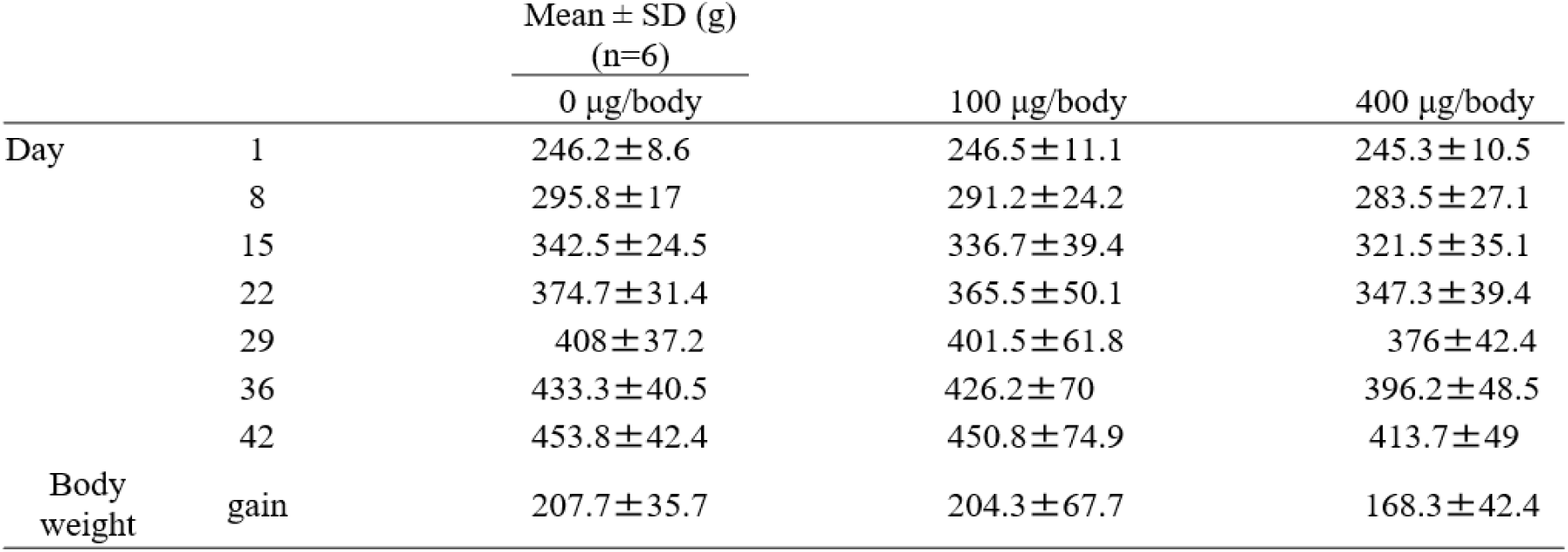
Effect of DNA vaccination on body weight in rats

**Table 3.** Effect of DNA vaccination on food consumption in rats

| | | Mean ± SD (g)<br>(n=6) | | |
| --- | --- | --- | --- | --- |
| | | 0 μg/body | 100 μg/body | 400 μg/body |
| Day | 8 | 26.35 ± 2.4 | 26.32 ± 3.84 | 24.37 ± 2.43 |
| | 15 | 28.23 ± 2.65 | 27.4 ± 5.11 | 25.38 ± 3.6 |
| | 22 | 27.6 ± 3.14 | 26.77 ± 4.61 | 25.38 ± 3.05 |
| | 29 | 27.45 ± 2.98 | 27 ± 4.8 | 25.12 ± 2.49 |
| | 36 | 27.45 ± 3.21 | 26.88 ± 4.84 | 25.15 ± 2.58 |
| | 42 | 26.28 ± 2.17 | 26.9 ± 4.98 | 24.62 ± 2.84 |

**Table 4.** Effect of DNA vaccination on Hematology in rats

| | | Mean ± SD | | |
| --- | --- | --- | --- | --- |
| | | 0 μg/body | 100 μg/body | 400 μg/body |
| RBC | $10^4/\mu\text{L}$ | 927.0±35.0 | 895.0±17.7 | 912.8±43.4 |
| HGB | g/dL | 16.65±0.66 | 16.52±0.23 | 16.70±0.73 |
| HCT | % | 45.47±2.09 | 45.37±1.08 | 46.12±2.23 |
| MCV | fL | 49.07±1.82 | 50.72±2.20 | 50.55±2.38 |
| MCH | pg | 17.98±0.54 | 18.47±0.58 | 18.30±0.71 |
| MCHC | g/dL | 36.63±0.36 | 36.43±0.51 | 36.22±0.57 |
| Reticulocyte | % | 3.268±0.261 | 3.120±0.557 | 3.288±0.439 |
| Platelet | $10^4/\mu\text{L}$ | 117.72±10.42 | 114.83±3.50 | 124.98±13.92 |
| WBC | $10^2/\mu\text{L}$ | 117.97±40.15 | 114.38±12.60 | 96.65±22.45 |
| Neutrophil | $10^2/\mu\text{L}$ | 23.22±11.07 | 15.18±2.39 | 13.08±4.70 |
| Lymphocyte | $10^2/\mu\text{L}$ | 89.22±30.14 | 93.23±10.75 | 78.87±17.66 |
| Monocyte | $10^2/\mu\text{L}$ | 3.97±1.24 | 4.70±1.28 | 3.17±1.35 |
| Eosinophil | $10^2/\mu\text{L}$ | 1.50±0.66 | 1.20±0.33 | 1.50±0.65 |
| Basophil | $10^2/\mu\text{L}$ | 0.07±0.08 | 0.07±0.05 | 0.03±0.05 |
| PT | sec | 18.50±1.68 | 17.88±1.88 | 19.35±1.13 |
| APTT | sec | 19.33±2.60 | 19.25±2.03 | 19.87±0.76 |

RBC: red blood cells, HGB: hemoglobine, HCT:hematocrit, MCV:mean colpuscular volume, MCH:mean colpuscular volume hemoglobine, MCHC:mean colpuscular volume hemoglobine concentration, WBC:white blood cells, PT:prothrombin time, APTT: activated partial thromboplastin time

**Table 5.**
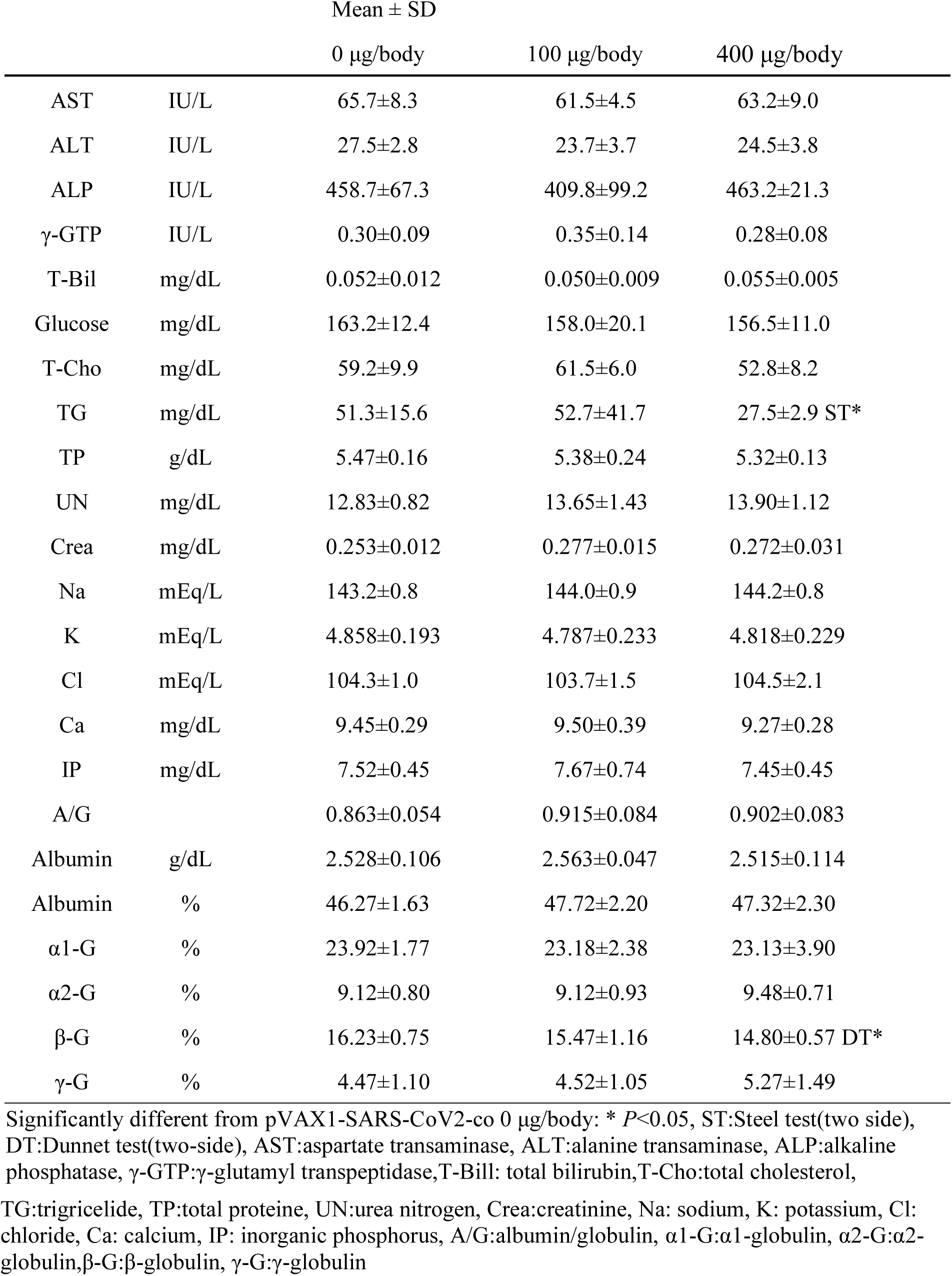
Effect of DNA vaccination on Blood biochemistry in rats

| | | Mean ± SD | | |
| --- | --- | --- | --- | --- |
| | | 0 μg/body | 100 μg/body | 400 μg/body |
| AST | IU/L | 65.7±8.3 | 61.5±4.5 | 63.2±9.0 |
| ALT | IU/L | 27.5±2.8 | 23.7±3.7 | 24.5±3.8 |
| ALP | IU/L | 458.7±67.3 | 409.8±99.2 | 463.2±21.3 |
| γ-GTP | IU/L | 0.30±0.09 | 0.35±0.14 | 0.28±0.08 |
| T-Bil | mg/dL | 0.052±0.012 | 0.050±0.009 | 0.055±0.005 |
| Glucose | mg/dL | 163.2±12.4 | 158.0±20.1 | 156.5±11.0 |
| T-Chol | mg/dL | 59.2±9.9 | 61.5±6.0 | 52.8±8.2 |
| TG | mg/dL | 51.3±15.6 | 52.7±41.7 | 27.5±2.9 ST* |
| TP | g/dL | 5.47±0.16 | 5.38±0.24 | 5.32±0.13 |
| UN | mg/dL | 12.83±0.82 | 13.65±1.43 | 13.90±1.12 |
| Crea | mg/dL | 0.253±0.012 | 0.277±0.015 | 0.272±0.031 |
| Na | mEq/L | 143.2±0.8 | 144.0±0.9 | 144.2±0.8 |
| K | mEq/L | 4.858±0.193 | 4.787±0.233 | 4.818±0.229 |
| Cl | mEq/L | 104.3±1.0 | 103.7±1.5 | 104.5±2.1 |
| Ca | mg/dL | 9.45±0.29 | 9.50±0.39 | 9.27±0.28 |
| IP | mg/dL | 7.52±0.45 | 7.67±0.74 | 7.45±0.45 |
| A/G | | 0.863±0.054 | 0.915±0.084 | 0.902±0.083 |
| Albumin | g/dL | 2.528±0.106 | 2.563±0.047 | 2.515±0.114 |
| Albumin | % | 46.27±1.63 | 47.72±2.20 | 47.32±2.30 |
| α1-G | % | 23.92±1.77 | 23.18±2.38 | 23.13±3.90 |
| α2-G | % | 9.12±0.80 | 9.12±0.93 | 9.48±0.71 |
| β-G | % | 16.23±0.75 | 15.47±1.16 | 14.80±0.57 DT* |
| γ-G | % | 4.47±1.10 | 4.52±1.05 | 5.27±1.49 |

Significantly different from pVAX1-SARS-CoV2-co 0 μg/body: \* $P$ <0.05, ST:Steel test(two side), DT:Dunnet test(two-side), AST:aspartate transaminase, ALT:alanine transaminase, ALP:alkaline phosphatase, $\gamma$ -GTP: $\gamma$ -glutamyl transpeptidase,T-Bil: total bilirubin,T-Chol:total cholesterol,
TG:triglyceride, TP:total protein, UN:urea nitrogen, Crea:creatinine, Na: sodium, K: potassium, Cl: chloride, Ca: calcium, IP: inorganic phosphorus, A/G:albumin/globulin, $\alpha$ 1-G: $\alpha$ 1-globulin, $\alpha$ 2-G: $\alpha$ 2-globulin, $\beta$ -G: β-globulin, $\gamma$ -G: $\gamma$ -globulin

**Table 6.**
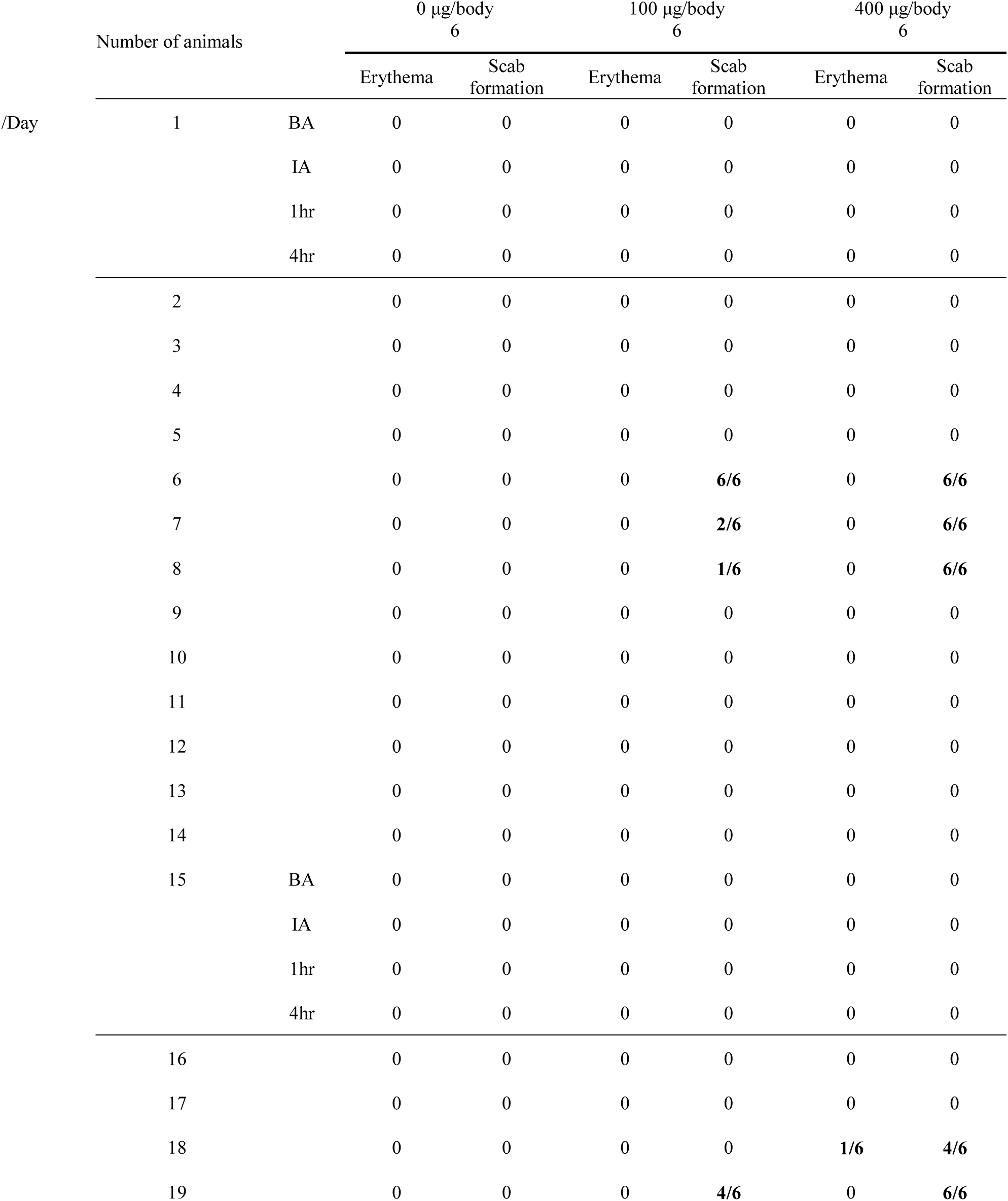
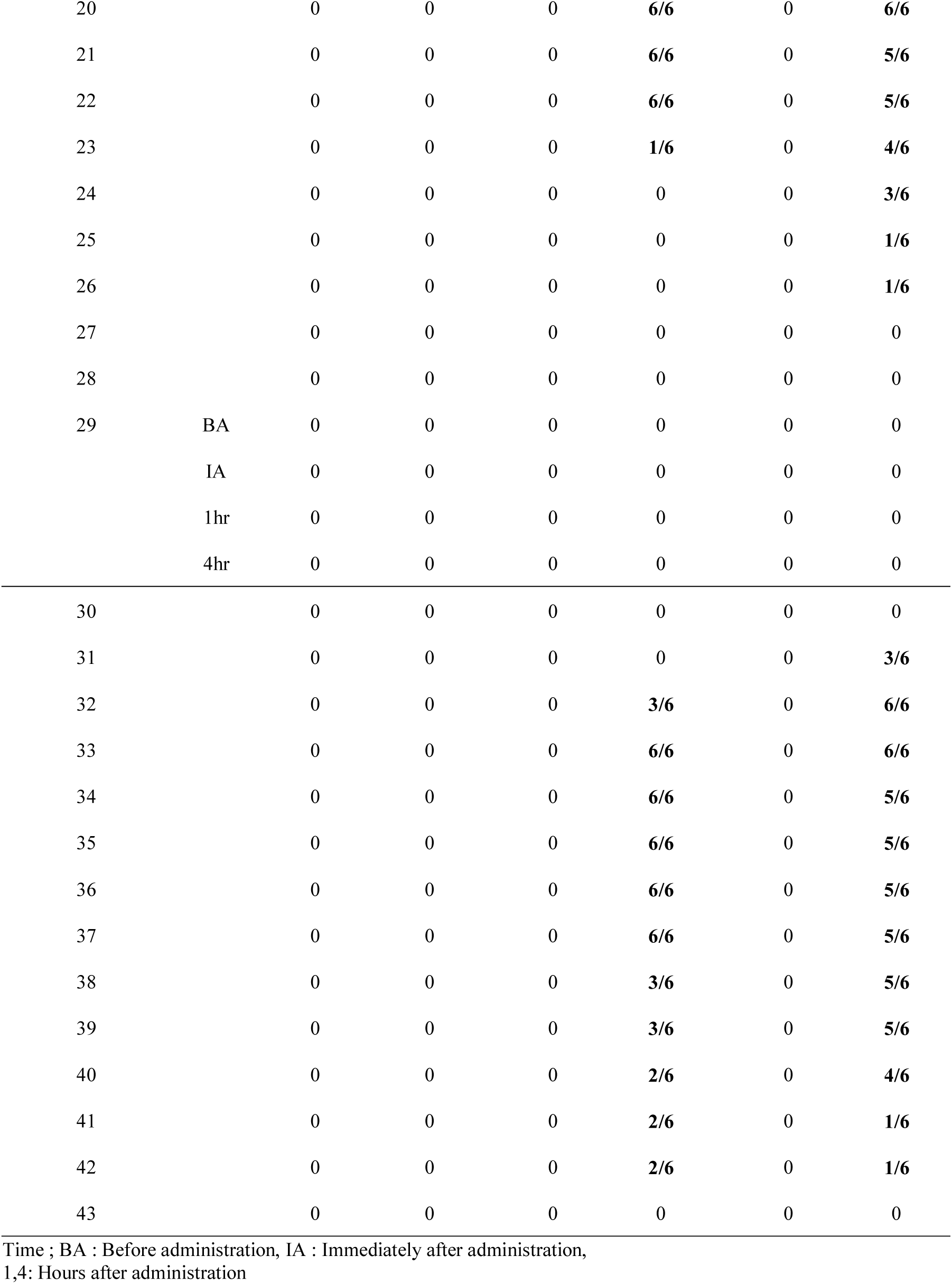
Clinical sign of DNA vaccination in rats

| Number of animals |  |  | 0 µg/body<br>6 |  | 100 µg/body<br>6 |  | 400 µg/body<br>6 |  |
| --- | --- | --- | --- | --- | --- | --- | --- | --- |
|  |  |  | Erythema | Scab<br>formation | Erythema | Scab<br>formation | Erythema | Scab<br>formation |
| /Day | 1 | BA | 0 | 0 | 0 | 0 | 0 | 0 |
|  |  | IA | 0 | 0 | 0 | 0 | 0 | 0 |
|  |  | 1hr | 0 | 0 | 0 | 0 | 0 | 0 |
|  |  | 4hr | 0 | 0 | 0 | 0 | 0 | 0 |
|  | 2 |  | 0 | 0 | 0 | 0 | 0 | 0 |
|  | 3 |  | 0 | 0 | 0 | 0 | 0 | 0 |
|  | 4 |  | 0 | 0 | 0 | 0 | 0 | 0 |
|  | 5 |  | 0 | 0 | 0 | 0 | 0 | 0 |
|  | 6 |  | 0 | 0 | 0 | **6/6** | 0 | **6/6** |
|  | 7 |  | 0 | 0 | 0 | **2/6** | 0 | **6/6** |
|  | 8 |  | 0 | 0 | 0 | **1/6** | 0 | **6/6** |
|  | 9 |  | 0 | 0 | 0 | 0 | 0 | 0 |
|  | 10 |  | 0 | 0 | 0 | 0 | 0 | 0 |
|  | 11 |  | 0 | 0 | 0 | 0 | 0 | 0 |
|  | 12 |  | 0 | 0 | 0 | 0 | 0 | 0 |
|  | 13 |  | 0 | 0 | 0 | 0 | 0 | 0 |
|  | 14 |  | 0 | 0 | 0 | 0 | 0 | 0 |
|  | 15 | BA | 0 | 0 | 0 | 0 | 0 | 0 |
|  |  | IA | 0 | 0 | 0 | 0 | 0 | 0 |
|  |  | 1hr | 0 | 0 | 0 | 0 | 0 | 0 |
|  |  | 4hr | 0 | 0 | 0 | 0 | 0 | 0 |
|  | 16 |  | 0 | 0 | 0 | 0 | 0 | 0 |
|  | 17 |  | 0 | 0 | 0 | 0 | 0 | 0 |
|  | 18 |  | 0 | 0 | 0 | 0 | **1/6** | **4/6** |
|  | 19 |  | 0 | 0 | 0 | **4/6** | 0 | **6/6** |

|  |  |  |  |  |  |  |  |
| --- | --- | --- | --- | --- | --- | --- | --- |
| 20 |  | 0 | 0 | 0 | **6/6** | 0 | **6/6** |
| 21 |  | 0 | 0 | 0 | **6/6** | 0 | **5/6** |
| 22 |  | 0 | 0 | 0 | **6/6** | 0 | **5/6** |
| 23 |  | 0 | 0 | 0 | **1/6** | 0 | **4/6** |
| 24 |  | 0 | 0 | 0 | 0 | 0 | **3/6** |
| 25 |  | 0 | 0 | 0 | 0 | 0 | **1/6** |
| 26 |  | 0 | 0 | 0 | 0 | 0 | **1/6** |
| 27 |  | 0 | 0 | 0 | 0 | 0 | 0 |
| 28 |  | 0 | 0 | 0 | 0 | 0 | 0 |
| 29 | BA | 0 | 0 | 0 | 0 | 0 | 0 |
|  | IA | 0 | 0 | 0 | 0 | 0 | 0 |
|  | 1hr | 0 | 0 | 0 | 0 | 0 | 0 |
|  | 4hr | 0 | 0 | 0 | 0 | 0 | 0 |
| 30 |  | 0 | 0 | 0 | 0 | 0 | 0 |
| 31 |  | 0 | 0 | 0 | 0 | 0 | **3/6** |
| 32 |  | 0 | 0 | 0 | **3/6** | 0 | **6/6** |
| 33 |  | 0 | 0 | 0 | **6/6** | 0 | **6/6** |
| 34 |  | 0 | 0 | 0 | **6/6** | 0 | **5/6** |
| 35 |  | 0 | 0 | 0 | **6/6** | 0 | **5/6** |
| 36 |  | 0 | 0 | 0 | **6/6** | 0 | **5/6** |
| 37 |  | 0 | 0 | 0 | **6/6** | 0 | **5/6** |
| 38 |  | 0 | 0 | 0 | **3/6** | 0 | **5/6** |
| 39 |  | 0 | 0 | 0 | **3/6** | 0 | **5/6** |
| 40 |  | 0 | 0 | 0 | **2/6** | 0 | **4/6** |
| 41 |  | 0 | 0 | 0 | **2/6** | 0 | **1/6** |
| 42 |  | 0 | 0 | 0 | **2/6** | 0 | **1/6** |
| 43 |  | 0 | 0 | 0 | 0 | 0 | 0 |

Time ; BA : Before administration, IA : Immediately after administration, 1,4: Hours after administration

Histopathological analysis of lymph nodes revealed slight to mild hyperplasia of follicular lymph nodes at the inguinal lymph node in 5 of 6 animals in the 400-µg injected group, slight hyperplasia of follicular was observed in 1 of 6 animals in the 100-µg injected group, and slight hyperplasia of follicular was observed at the axillary lymph node of one of six animals in the 400-µg injected group. Furthermore, hyperplasia of plasma cells was observed only in the 400-µg injected group. These abnormalities were not observed in the PBS-injected group; thus, this hyperplasia might be an immune response against pVAX1-SARS-CoV2-co injection. Sperm reduction in the epididymis, interstitial inflammation of the prostate, or sperm granuloma were observed; however, these were also observed in the PBS-injected group (Table7).

## Discussion

In the recent COVID-19 pandemic, efficient and safe DNA vaccine development is an important issue. Thus, we tested the immunogenicity and safety of the combination of human codon-optimized new COVID-19 DNA vaccine candidate (pVAX1-SARS-CoV2-co)^4^ and PJI^7^. Development of vaccines against SARS-CoV-2 is the priority in many pharmaceutical companies and academic organizations all over the world. A variety of SARS-CoV-2 vaccines, such as protein subunit^12^, replicating or non-replicating viral vector^13-15^, inactivated virus^16^, live attenuated virus^17-18^, virus-like particle (VLP)^19-20^, RNA-^21-22^ or DNA-based^6, 23-24^ vaccines, are currently under development. DNA vaccines have several advantages compared to traditional vaccines, such as protein subunits, peptides, and living/inactivated pathogens. Compared to a protein- or peptide-based vaccines or, living/inactivated virus vaccines, DNA vaccine does not require the cultivation of the target pathogen and can be produced on an industrial scale^25^. Furthermore, rapid adaptation can be applied when mutations occur in the target DNA/RNA sequence. Moreover, the vaccine administration is another important factor in enhancing efficacy. In recent years, the major vaccination method has been intramuscular administration using a needle syringe. Intradermal administration, however, is also a promising vaccination method because dermal tissue is an easily accessible immune-rich environment that can induce both humoral and cellular immunity^26^. Tebas *et al*. reported that intradermal Ebola GP DNA vaccine administration demonstrates humoral and cellular immunogenicity advantages compared with intramuscular administration^27^.

When 100 or 400 µg of pVAX1-SARS-CoV2-co was injected into the rat, SARS-CoV-2 spike protein (S1+S2) antibody was detected (Fig 4A). Statistically significant differences were observed between the 100- and 400-µg pVAX1-SARS-CoV2-co-injected group after 4 and 6 weeks. This meant that the dose-dependent anti-SARS-CoV-2 spike protein (S1+S2) antibody production was induced as described in our previous study using the OVA expression plasmid DNA^4^. Interestingly, the antibody was detected in the 2-week sample for both doses. This meant that the antibody release was induced quickly after the first injection. The potential for rapid antibody production seemed likely for the vaccination method. In addition, antibody titers were increased eight times in the low-dose group and 37 times in the high-dose group after the second injection, and 2-3 times after the third injection. This indicated the importance of the second injection as an immune booster. Moreover, the number of IFN-γ-producing cells were increased in the pVAX1-SARS-CoV2-co-injected group, indicating that cellular immunity was elicited via pVAX1-SARS-CoV2-co intradermal injection. To prevent and cure the viral infection, the elicitation of cellular immunity is important. Thus, the combination of pVAX1-SARS-CoV2-co and intradermal injection via PJI is a promising method against SARS-CoV-2.

In the safety test, no animals had died by the time of necropsy and there was no significant difference in body weight, food consumption, urinalysis, organ weight, and hematology between PBS- and pVAX1-SARS-CoV2-co-injected groups. As shown in Table 5, blood biochemistry revealed significantly lower levels of triglyceride and β-globulin in the 400-µg µg group than in the PBS group. However, these changes were all within the variation of historical data of normal animals. These results suggested that pVAX1-SARS-CoV2-co injection at these doses did not cause systemic toxicity in rats. Scab formulation was observed in the injected region, and the duration between the injection and scab formation became shorter with the vaccination time. Also, the duration of scab in the high-dose group was shorter than that in the low-dose group (Table 6). These results suggested that scab formation was related to the immune response caused by the pVAX1-SARS-CoV2-co injection. However, all scabs disappeared within 14 days of injection. Slight to mild hyperplasia of follicular or plasma cells was observed in the inguinal lymph node of 5 in 6 animals and the axillary lymph node of 1 in 6 animals in the high-dose group, whereas it was observed in the inguinal lymph node of 1 in 6 animals in the low-dose group (Table 7). The immune response of the high-dose group was higher than that in the low-dose group. Thus, hyperplasia was thought to be an immune response against the antigen invasion, which was not toxic. Sperm reduction in the epididymis, interstitial inflammation of the prostate, or sperm granuloma were observed; however, these were observed in the PBS-injected group as well. Thus, these observations did not indicate any toxic signs.

**Table 7.**
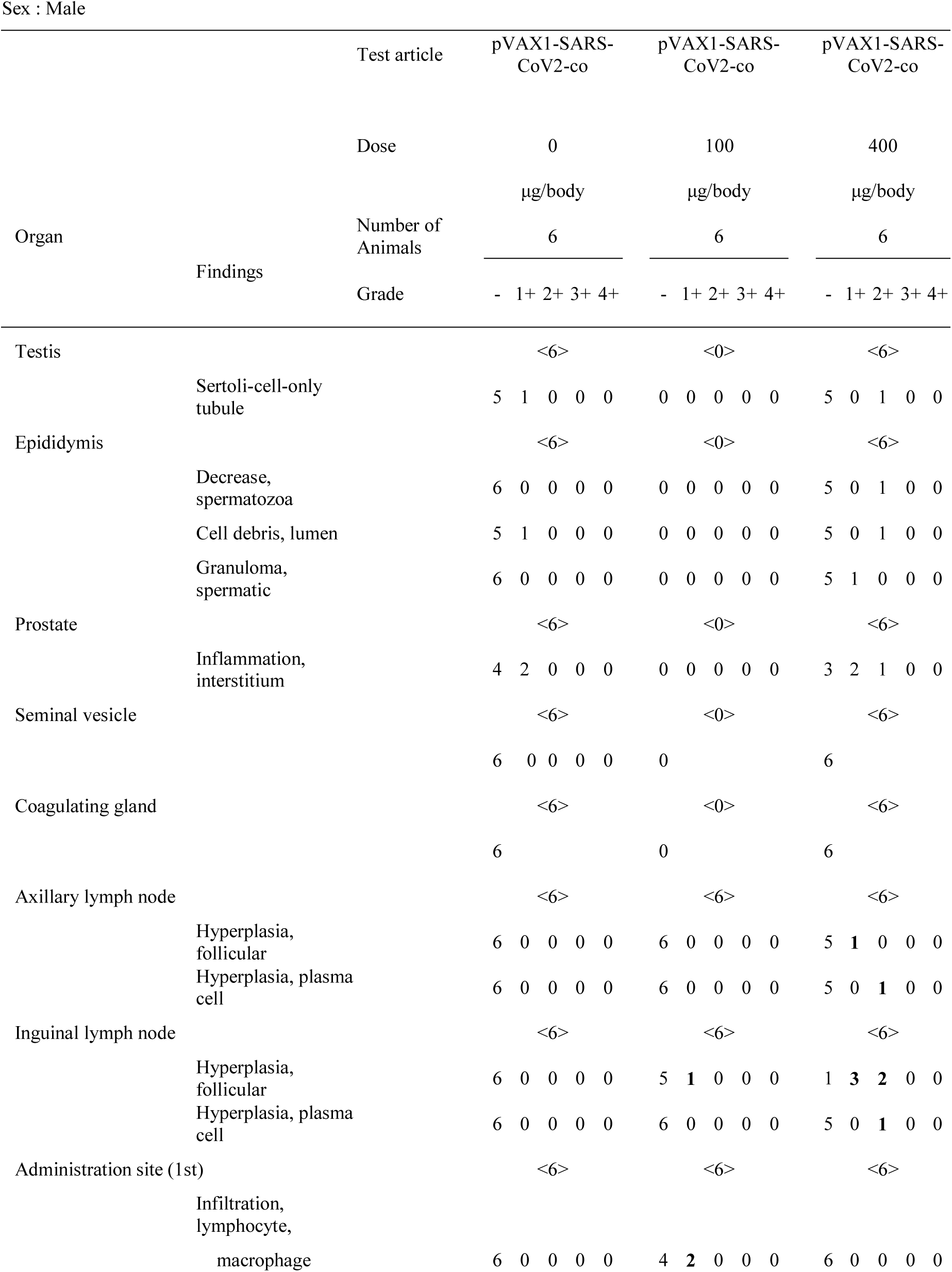
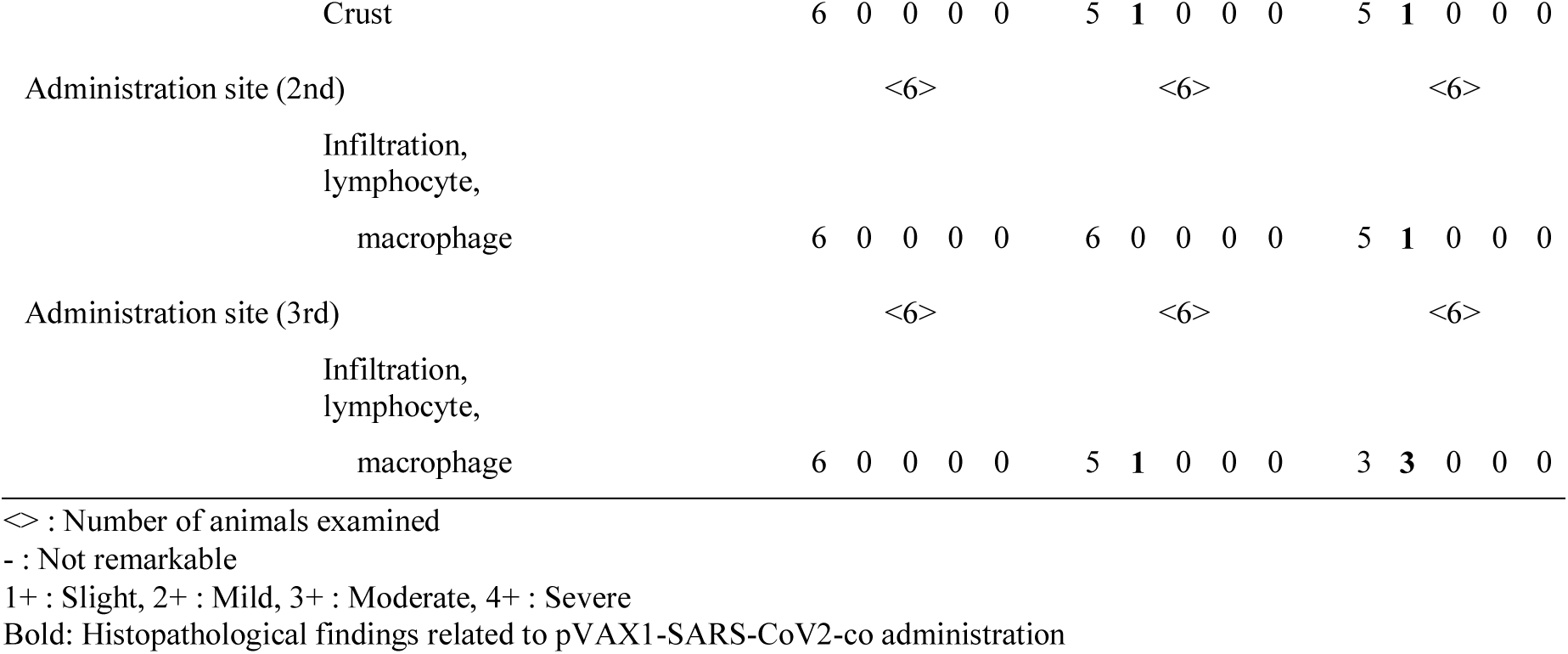
Histopathological findings

| Organ | Findings | Test article | pVAX1-SARS-CoV2-co | pVAX1-SARS-CoV2-co | pVAX1-SARS-CoV2-co |
| --- | --- | --- | --- | --- | --- |
|  |  | Dose | 0 | 100 | 400 |
|  |  |  | µg/body | µg/body | µg/body |
|  |  | Number of Animals | 6 | 6 | 6 |
|  |  | Grade | - 1+ 2+ 3+ 4+ | - 1+ 2+ 3+ 4+ | - 1+ 2+ 3+ 4+ |
| Testis |  |  | <6> | <0> | <6> |
|  | Sertoli-cell-only tubule |  | 5 1 0 0 0 | 0 0 0 0 0 | 5 0 1 0 0 |
| Epididymis |  |  | <6> | <0> | <6> |
|  | Decrease, spermatozoa |  | 6 0 0 0 0 | 0 0 0 0 0 | 5 0 1 0 0 |
|  | Cell debris, lumen |  | 5 1 0 0 0 | 0 0 0 0 0 | 5 0 1 0 0 |
|  | Granuloma, spermatic |  | 6 0 0 0 0 | 0 0 0 0 0 | 5 1 0 0 0 |
| Prostate |  |  | <6> | <0> | <6> |
|  | Inflammation, interstitium |  | 4 2 0 0 0 | 0 0 0 0 0 | 3 2 1 0 0 |
| Seminal vesicle |  |  | <6> | <0> | <6> |
|  |  |  | 6 0 0 0 0 | 0 | 6 |
| Coagulating gland |  |  | <6> | <0> | <6> |
|  |  |  | 6 | 0 | 6 |
| Axillary lymph node |  |  | <6> | <6> | <6> |
|  | Hyperplasia, follicular |  | 6 0 0 0 0 | 6 0 0 0 0 | 5 1 0 0 0 |
|  | Hyperplasia, plasma cell |  | 6 0 0 0 0 | 6 0 0 0 0 | 5 0 1 0 0 |
| Inguinal lymph node |  |  | <6> | <6> | <6> |
|  | Hyperplasia, follicular |  | 6 0 0 0 0 | 5 1 0 0 0 | 1 3 2 0 0 |
|  | Hyperplasia, plasma cell |  | 6 0 0 0 0 | 6 0 0 0 0 | 5 0 1 0 0 |
| Administration site (1st) |  |  | <6> | <6> | <6> |
|  | Infiltration, lymphocyte, macrophage |  | 6 0 0 0 0 | 4 2 0 0 0 | 6 0 0 0 0 |

|  |  |  |  |  |  |  |  |  |  |  |  |  |  |  |  |
| --- | --- | --- | --- | --- | --- | --- | --- | --- | --- | --- | --- | --- | --- | --- | --- |
| Crust | 6 | 0 | 0 | 0 | 0 | 5 | **1** | 0 | 0 | 0 | 5 | **1** | 0 | 0 | 0 |
| Administration site (2nd) | <6> |  |  |  |  | <6> |  |  |  |  | <6> |  |  |  |  |
| Infiltration,<br>lymphocyte,<br>macrophage | 6 | 0 | 0 | 0 | 0 | 6 | 0 | 0 | 0 | 0 | 5 | **1** | 0 | 0 | 0 |
| Administration site (3rd) | <6> |  |  |  |  | <6> |  |  |  |  | <6> |  |  |  |  |
| Infiltration,<br>lymphocyte,<br>macrophage | 6 | 0 | 0 | 0 | 0 | 5 | **1** | 0 | 0 | 0 | 3 | **3** | 0 | 0 | 0 |

<> : Number of animals examined
- : Not remarkable
1+ : Slight, 2+ : Mild, 3+ : Moderate, 4+ : Severe
**Bold:** Histopathological findings related to pVAX1-SARS-CoV2-co administration

We evaluated the safety of intradermal vaccination of human codon-optimized COVID-19 DNA vaccine candidate (pVAX1-SARS-CoV2-co). Intradermal PJI injection of pVAX1-SARS-CoV2-co did not show any systemic toxicity, whereas erythema and scab formation were observed at skin injection sites. However, the visual observation and histopathological analysis indicated that these abnormalities were not permanent and expected to fully recover in time. No other toxic signs were observed. Thus, these results indicate that intradermal administration of pVAX1-SARS-CoV2-co is a safe and promising method to induce potent antibodies against SARS-CoV-2.

## Materials and Methods

### Plasmids

pVAX1-SARS-CoV2-co was designed by Takara, which encoded a highly optimized DNA sequence encoding the SARS-CoV-2 Spike glycoprotein. The plasmid DNA was amplified and purified using a GIGA prep (Qiagen, USA).

### Animals

The rat study was performed using 7-week-old male Crl:CD (SD) rats (Charles River Laboratories Japan Inc., Kanagawa, Japan). The mouse study was performed using 8-10 weeks old C57BL/6NCrSlc mice (C57BL/6N) (Japan SLC,Inc.). All animals were maintained under controlled conditions (temperature: 21.0-24.5 °C, humidity: 45 ± 15%, ventilation: 8–15 times/h, light/dark cycle: 12 h) in a pathogen-free room. Animals received free access to food and water and were handled according to the approved protocols of the Animal Committee of Osaka University (Suita, Osaka, Japan) as No.28-021-028, the Ethics Committee for Animal Experiments of the Safety Research Institute for Chemical Compounds Co. Ltd. (Sapporo, Hokkaido, Japan) as No. AN20200617-03 and the Animal Committee of KAC Co. Ltd. (Kusatsu, Shiga, Japan) as No. 20–0508. All mice experiments were conducted with the approval of the Animal Research Committee of Research Institute for Microbial Disease in Osaka University (Suita, Osaka, JAPAN).

### Immunogenicity test

Two groups of six animals received pVAX1-SARS-CoV2-co injection in their flunk areas on both sides every 2 weeks 3 times. For each injection, animals received 30μL plasmid solution injection at two sites on each side. The injected areas were changed for every injection to prevent repeated injections at the same site. The sera were collected every 2 weeks under anesthesia until 12 weeks for the first and second groups for the ELISpot assay.

### Safety evaluation

Animals were divided into three groups of six animals, and animals received a pVAX1-SARS-CoV2-co injection in their flunk areas on both sides every 2 weeks for three times. For each injection, the first group received 50 μL of PBS solution at four sites, the second group received 50 μL of plasmid solution at one site, and the third group received 50 μL of plasmid solution at four sites; thus, each rat received 0, 100, and 400 μg of DNA vaccination in each group. The injected areas were changed after every injection to prevent repeated injections at the same skin site. To assess the immune response, sera were collected every 2 weeks under anesthesia until 6 weeks and on day 43, rats were sacrificed. In both experiments, the PJI, Actranza™ lab. for Rat was employed for intradermal injection, supplied by Daicel Corporation.

During the treatment period, animals were observed for general signs and observed for skin reaction of the injected region every day. Body weight and food consumption were measured once a week. On the day of necropsy, blood for hematology, blood biochemistry, and immune-response assays was collected under isoflurane anesthesia. Urine was collected for urinalysis. After euthanasia and whole blood sampling, the skin region, inguinal lymph node, reproductive organs, and other organs that were found to be abnormal upon visual observation were extracted for microscopic examination. All injected skin sites were excised to assess the recovery of the injection site. The sampled organs were fixed with 10% neutral buffered formalin solution. Thin paraffin sections were prepared and stained with hematoxylin-eosin and subjected to histopathological analysis.

### ELISA

Precisely 1 μg/ml of recombinant 2019-nCoV spike protein (S1+S2) (ECD, His tag) (BLPSN-0986P, BETA Life Sciences, Fairfield, NJ, USA) was immobilized on a 96-well plate (442404, MAXISORP F96 NUNC Immuno-plate, Thermo Fisher Scientific, Roskilde, Denmark) at 4 °C overnight. After discarding the recombinant protein solution, the plate was incubated with 5% skim milk in PBS (blocking buffer) for 2 h at room temperature. After blocking, diluted rat sera (50-, 250-, 1250-, 6250-, 3250- and 156250-fold dilutions for S1+S2 or 10-, 50-, 250-, 1250-, 6250-, 31250-, and 156250-fold dilutions for RBD) or diluted mouse sera (50-, 250-, 1250-, 6250-, 3250-, and 156250-fold dilutions for S1+S2 or 50-, 250-, 1250-, 6250-, and 31250-fold dilutions for RBD) were incubated at 4°C overnight. All wells were washed with PBST (200 μl/well) seven times, and incubated with 1:1000 diluted Amersham ECL anti-rat IgG horseradish peroxidase-linked species-specific whole antibody (NA935, GE Healthcare UK Limited, UK) or Amersham ECL anti-mouse IgG horseradish peroxidase-linked species-specific whole antibody (NA931, GE Healthcare UK Limited, UK) in the blocking buffer for 3 h at room temperature. After washing with 50 μl/well of PBST four times, 50 μl/well of 3,3’-5,5’-tetramethylbenzidine (TMB) Liquid Substrate System for ELISA (T040, Sigma-Aldrich Co. LLC., St. Louis, MO) was added to the plate. After 30 min of incubation, 50 μl/well of 0.9 N H2SO4 solution was added and OD450nm was measured using an iMark Microplate Reader (1681135J, Bio Rad, Hercules, CA). The values of the half-maximal antibody titers of serum samples were calculated from the highest absorbance in the dilution range using Prism 8 (Graph Pad Software Inc. San Diego, CA 92018, USA).

### ELISpot assays

For splenocyte preparations, rat spleens were harvested from the vaccinated rats after 6 weeks. Splenocytes were passed through a 70-μm cell strainer in RPMI 1640 medium (Nacalai Tesque Inc., Kyoto, Japan) supplemented with 0.1 mg/ml penicillin-streptomycin (Nacalai Tesque Inc.). Residual erythrocytes were lysed using hemolysis buffer (Immuno-Biological Laboratories Co., Ltd., Gunma, Japan) for 5 min and washed with PBS. Splenocytes were subsequently cultured in complete RPMI 1640 medium supplemented with 10% fetal bovine serum (FBS; GE Healthcare Life Sciences, Logan, UT), 0.1 mg/ml penicillin-streptomycin, and 50 μM β-mercaptoethanol (Nacalai Tesque Inc.) and incubated at 37 °C in a humidified atmosphere of 5% CO2. Rat IFN-γ and IL-4 ELISpot assays (CT079 and CT081, U-CyTech biosciences, Utrecht, Netherlands) were performed according to the manufacturer’s instructions. Rat splenocytes (2 × 10^5^) were plated in duplicate in 96-well PVDF plates (Millipore, Billerica, MA) and stimulated with 5 μg/ml of recombinant 2019-nCoV Spike protein (S1+S2 ECD, His tag; Beta Lifescience, Fairfield, NJ) for 48 h. The resultant stained well membranes were scanned, and the number of spot-forming cells was counted.

### IgG subclass analysis

To analyze the immunoglobulin subclass profile in sera of vaccinated rats, standardized ELISA protocols were performed. Briefly, 96-well plates were coated with 10 µg/ml SARS-CoV-2 SV2 recombinant protein at 4 °C overnight. Serum samples were incubated in appropriate wells at 4 °C overnight. After serum incubation, the serum samples were removed and IgG subtypes were detected using the following HRP-conjugated antibodies for 3 h at room temperature; anti-IgG1 H&L (ab106753; Abcam), anti-IgG2a H&L (ab106783; Abcam). anti-IgG2b H&L (ab106750; Abcam), and anti-IgG2c (3075–05, Southern Biotech). After HRP-conjugated antibody detection, plates were developed using POD (T0440; Sigma) and stopping buffer (9562606; Nacalai Tesque) and the absorbance was detected at 450 nm (iMark; Bio rad).

### ACE2 binding inhibition assay

To analyze the binding inhibition of S1+S2 and ACE2 by neutralizing antibodies in the immunized rat and mouse serum, 96-well plates were coated with human ACE2 recombinant protein (1μg/ml, mFc tag; #83986, Cell Signaling Technology), and then blocked using PBS containing 5% skim milk for 2 h at room temperature. After blocking, pre-incubated samples of diluted rat or mouse serum with recombinant S1+S2 protein (2 μg/ml, His tag; Beta Lifescience) was added to the coated wells and incubated at 4 °C overnight. Wells were washed with PBST and then incubated with anti-His antibody conjugated with HRP (GTX21187, GeneTex, Inc., Irvine, CA) for 2 h at room temperature. After washing with PBST, the peroxidase chromogenic substrate TMB (Sigma-Aldrich) was added to the wells and incubated for 30 min at room temperature. The reaction was stopped by adding 0.5 N sulfuric acid. The absorbance at 450 nm was measured using a microplate reader (Bio rad). Percentage inhibition of immunized blood serum at various time points was calculated and plotted using GraphPad Prism software, from which the neutralization titer at ID75 was derived.

### Pseudovirus neutralization assay for SARS-CoV-2

The neutralizing activity of vaccine-induced antibodies was analyzed using pseudotyped VSVs, as previously described^10^. Briefly, Vero E6 cells stably expressing TMPRSS2 were seeded on 96-well plates and incubated at 37 °C for 24 h. Pseudoviruses were incubated with a series of dilutions of inactivated rat serum for 1 h at 37 °C and then added to Vero E6 cells. Exactly 24 h after infection, the cells were lysed with cell culture lysis reagent (Promega), and luciferase activity was measured using a Centro XS3 LB 960 (Berthold).

### SARS-CoV-2 infection in mice that received pVAX1-SARS-CoV2-co plasmids

Female mice intradermally received pVAX1-SARS-CoV-2-co plasmids. For the ID injection, animals received 160 µg of plasmid at two sites on each flank using PJI for mouse of Actranza™ lab. Animals received 20μl (40μg) pVAX-SARS-CoV-2-co plasmid at two sites on each side, totally 160μg of plasmid for each mouse, using PJI. pVAX1-SARS-CoV-2-co plasmids were administered on days 0 and 14. Sixteen weeks after the first administration, sera were collected from immunized mice and tested for antibody induction against 2019-nCoV spike protein (S1+S2) and the RBD region of the spike protein, as well as for the inhibition of binding to the recombinant hACE2 protein.

The mouse-adapted SARS-CoV-2 virus (HuDPKng19-020 strain-Q498Y/P499T) was generated in vitro using reverse genetics^11, 28^. The infectious titers in the culture supernatants were determined analyzing the 50% tissue culture infective doses (TCID50). Mice were intranasally infected with 2.0 × 10^5^ TCID50 mouse-adapted SARS-CoV-2 virus. Two days post infection, lungs were isolated and 0.02 g of the tissue was homogenized using 100 µl of PBS. The homogenate solution was serially diluted 10 folds in DMEM containing 2% FBS and loaded on VeroE6/TMPRSS2 cells to determine the value.

### Statistics

Body weight, food consumption, hematology and blood biochemistry, urinalysis, and organ weight were analyzed using MiTox (Mitsui Toxicological Data Processing System, Mitsui E&S Systems Research Inc., Tokyo, Japan). The statistical analysis for antibody titers and ELISpot was performed using BellCurve for Excel (Social Survey Research Information Ltd., Tokyo, Japan). A *p*-value less than 0.05 was considered statistically significant.

## Acknowledgments

This research was supported by the Japan Agency for Medical Research and Development (AMED) under grant number JP20nk0101602.

---

**Supplementary Figure 1.**
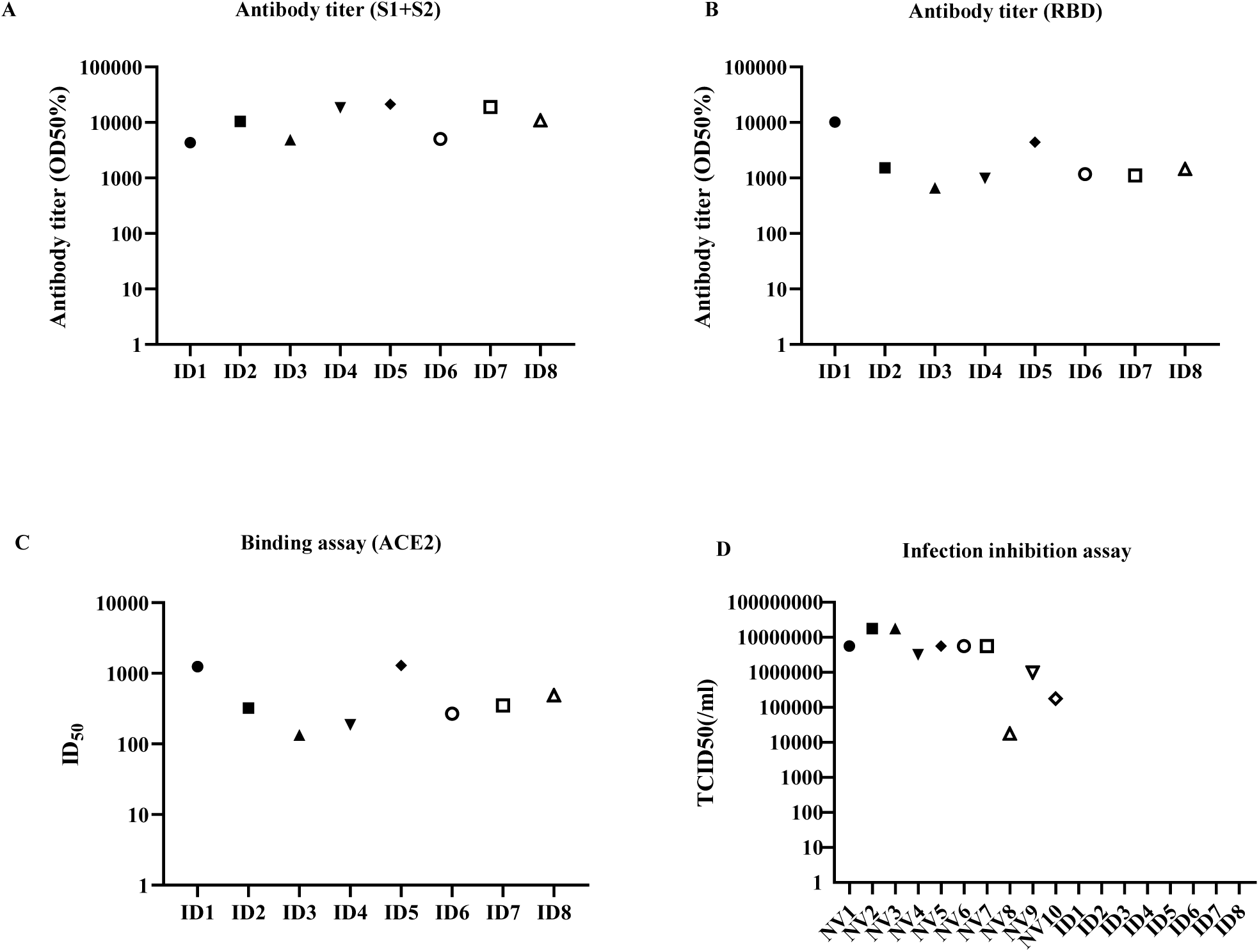
Individual values of pre-viral challenge antibody titers of ID and NV mice for recombinant S1+S2 (A), RBD (B) and neutralization titer (inhibitory dose at 50% neutralization; ID50) of ACE2-S1+S2 binding inhibition (C), as well as TCID50 values in the mouse lung tissues after virus challenge as described in Fig. 5

